# Simultaneous CO_2_ and CO methanation using microbes

**DOI:** 10.1101/326561

**Authors:** Kohlmayer Matthias, Huber Robert, Brotsack Raimund, Mayer Wolfgang

**Author notes:** Address correspondence to Kohlmayer Matthias,.

## Abstract

In this study, we developed a method for simultaneous bio-methanation of CO_2_ and CO with H_2_ in a single bioreactor using a combination of carboxydotrophic bacteria and methanogenic archaea for industrial applications. Methanogenic archaea generally use H_2_ and CO_2_ to produce methane, whereas very few methanogenic archaea methanize CO, and these grow slowly and consequently produce low reactant gas turnover rates. Thus, to achieve fast and simultaneous transformation of CO and CO_2_, we identified a combination of carboxydotrophic and hydrogenogenic bacteria and methanogenic archaea that can produce H_2_ and CO_2_ from CO, and then methanize CO_2_ and H_2_. The present screening experiments identified carboxydotrophic bacteria and methanogenic archaea that can cohabitate at the same thermophilic temperature and pH ranges and in the same growth medium. In these experiments, combinations of *Carboxydocella thermautotrophica* (DSM 12326), *Carboxydocella sporoproducens* (DSM 16521), and three thermophilic rod-shaped methanogenic archaeal cultures from MicroPyros GmbH formed unique microbial co-cultures that transformed CO_2_, H_2_, and CO to methane. The successful combination of these microbes could be used to gasify biowastes, such as sewage sludge, as alternative sources of hydrogen for microbial power-to-gas processes. Accordingly, gasification under these conditions produced H_2_-rich gas containing CO_2_ and CO, theoretically allowing various types of biowastes to be converted to biomethane, which is CO_2_-neutral, storable, and widely applicable as an energy source.

## IMPORTANCE

In this study, we hypothesized that the simultaneous bio-methanation of CO_2_ and CO with H_2_ in a single bioreactor can support the Power-to-Gas technology, a storage technology for renewable energies. We formed a novel co-culture tool that efficiently achieved the fast and simultaneous transformation of CO and CO_2_. That novel co-culture consists of *Caroxydocella thermautotrophica* (DSM 12326), *C. sporoproducens* (DSM 16521), and three thermophilic rod-shaped methanogenic archaeal cultures from MicroPyros GmbH.

## Introduction

We hypothesized that the efficient catalysis of the methanation of CO_2_ and CO in a single bioreactor can aid in promoting the expansion of renewable energy sources to slow climate change. For the efficient catalysis, we have developed a combination of carboxydotrophic bacteria and methanogenic archaea.

Power-to-Gas (PtG) storage technologies have been developed to support the use of renewable energies, and these use electrical power from renewable sources to split water and generate hydrogen and oxygen. Hydrogen from such hydrolysis reactions can be stored as methane following reactions with CO_2_ as follows: 4 H_2_ + CO_2_–> CH_4_ + 2 H_2_O. Hence, catalysis of such reactions by methanogenic archaea will constitute a microbial PtG technology.

The workload and profitability of PtG technologies can be increased using alternative sources of hydrogen when electrical power for hydrogen production is expensive. To this end, gasification of sewerage sludge may provide an adjustable, ecological, and economically rational alternative H_2_ source for microbial PtG processes (1), (2). The ensuing gasification produces H_2_-rich gas that also contains CO_2_ and CO, and could be used directly as a gas mixture for the PtG processes, pending on the methanation of CO.

In contrast with chemical-catalyzed synthesis of methane from H_2_ and CO_2_ using the so-called Sabatier process, microorganism biocatalysts can adapt to gas pollution, are robust to fluctuations of gas contents, and can work at lower temperatures and pressures (3). Multiple methanogenic archaea have been shown to methanize CO_2_ and H_2_, although very few reportedly methanize CO (4), (5), (6), (7), and these grow slowly and produce low rates of reactant gas turnover (8). However, efficient CO-oxidizing, H_2_-producing bacteria have been identified, and these use CO as their only source of carbon and energy (9).

Klasson *et al.* (10) previously reported the use of a single co-culture comprising *Rhodospirillum rubrum* to convert CO and H_2_ to methane and *Methanobacterium formicicum* and *Methanosarcina barkeri* to convert H_2_ and CO_2_ to methane at 34°C. Herein, we identified microbes that could simultaneously methanize CO_2_, H_2_, and CO at thermophilic growth temperatures. We then devised a novel combination of CO-utilizing and H_2_-producing bacteria (CO + H_2_O–> CO_2_ + H_2_) with methanogenic archaea (CO_2_ + 4 H_2_–> CH_4_ + 2 H_2_O) and showed optimal growth at around 63°C.

Our data demonstrate a highly efficient microbial co-culture that transforms CO_2_, H_2_, and CO to methane. This co-culture comprised a combination of *Carboxydocella thermautotrophica* (DSM 12326), *Carboxydocella sporoproducens* (DSM 16521), and three methanogenic archaea cultures from MicroPyros GmbH. The present innovation may provide the basis for gasification of sewage sludge as an alternative source of hydrogen for industrial scale microbial PtG processes.

## Materials and Methods

Based on the methods reported by Hofbauer *et al.* (11), we used the synthetic steam gasifier and synthetic air gasifier gas compositions shown in Table 1. Gases were purchased from Tyczka Industrie-Gase of Mannheim, Germany.

**Table 1.**
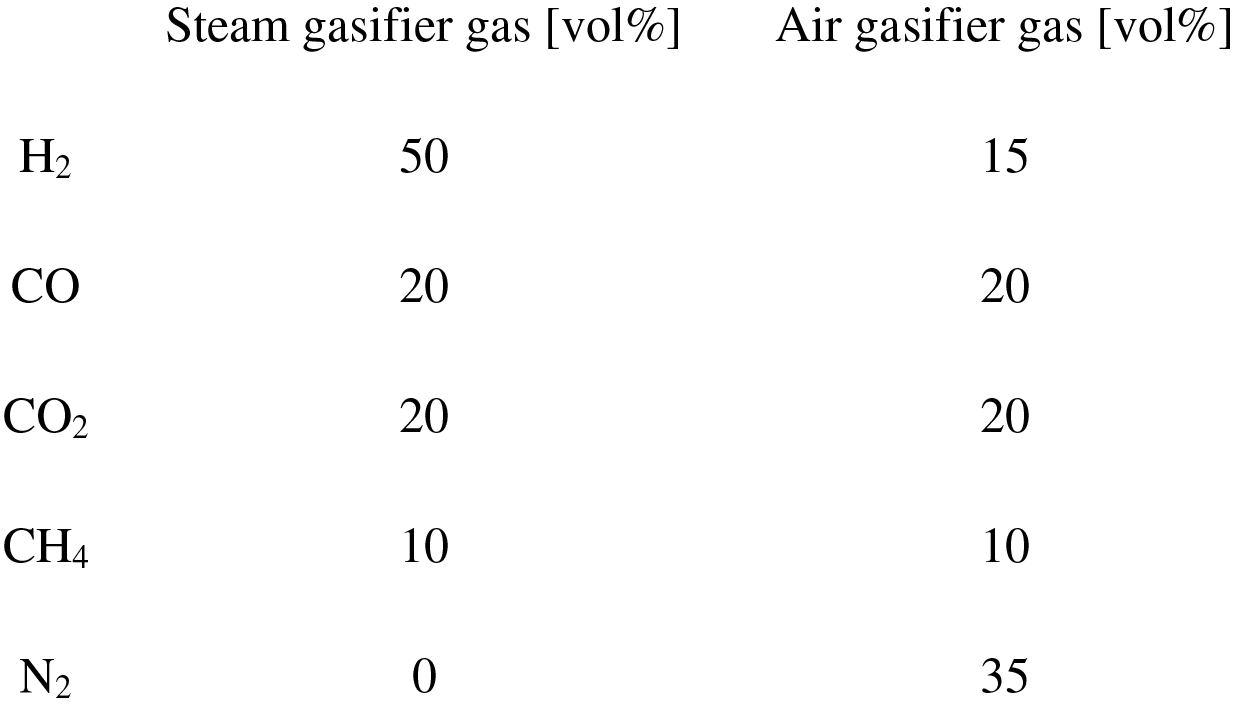
*Gas concentrations of steam gasifier and air gasifier gases*

|  | Steam gasifier gas [vol%] | Air gasifier gas [vol%] |
| --- | --- | --- |
| H<sub>2</sub> | 50 | 15 |
| CO | 20 | 20 |
| CO<sub>2</sub> | 20 | 20 |
| CH<sub>4</sub> | 10 | 10 |
| N<sub>2</sub> | 0 | 35 |

The present co-cultures were established as combinations of *Carboxydocella thermautotrophica* (DSM 12326; 12), *Carboxydocella sporoproducens* (DSM 16521; 13; so-called Carbos) from Deutsche Sammlung für Mikroorganismen und Zellkulturen of Brunswick, Germany, and a cell mixture of three thermophilic rod-shaped methanogenic archaeal cultures (TRMC) that were kindly supplied by MicroPyros GmbH of Straubing, Germany. TRMC are autotrophic isolates from the biogas plant of Zweckverband Abfallwirtschaft of Straubing, Germany and were selected in screens for reproducible growth in newly designed medium (see below) with synthetic steam gasifier gas and synthetic air gasifier gas. The present isolates showed the best and most reproducible growth among all strains tested and were selected from the MicroPyros culture collection. *Carboxydocella sporoproducens* cells grow more slowly than *Carboxydocella thermautotrophica* cells but can form stable spores that survive periods of CO deprivation (12), (13). To ensure sustained supply, cultures were transferred into fresh media weekly (5% inoculation) and were incubated at 63°C for 24 h. All experiments were performed in triplicate for at least five transfers.

Growth media were prepared as described by Zhao *et al.* (14), except that the expensive vitamin and yeast extract supplements were omitted for future industrial scale-up. The final media contained (g/L) KCl (0.33), MgCl_2_×6H_2_O (0.102), CaCl_2_O×2H_2_O(0.015), NH_4_Cl (0.33), K_2_HPO_4_ (0.14), and NaHCO_3_ (0.42) in demineralized water. A trace element solution (10 mL/L) and a selenite-tungstate solution (0.1 mL/L) were added as described for DSMZ medium 14 and DSMZ medium 385.

Oxygen was expelled from media solutions using the argon method described by Hungate (15) with minor modifications. Subsequently, 0.7 g/L of Na_2_S×9H_2_O was added, and media were introduced to anaerobic chambers and were dispensed into 28 mL serum tubes at 5 mL per tube. Tubes were then sealed with rubber stoppers and were pressurized to 3 bar absolute using the desired gas phase. Gas phases in glass tubes were then exchanged by degassing and gassing three times. Pressurized tubes were then autoclaved at 121°C for 15 min (15), (16) and the pH of the autoclaved medium was about 6.5. Culture tubes were incubated almost horizontally to maximize contact areas of gas and liquid phases, and gases entered liquid phases containing microorganisms solely by diffusion. Samples of cultures were taken using syringes (Omnifix-F^®^), and numbers of cells were counted using fluorescence microscopy (Olympus BX53F) with a Thoma counting chamber. Numbers of cells in TRMC were determined by counting fluorescent cells, and pH was determined using pH sticks (pH-Fix; 4.5–10.0, Roth). Pressure levels in culture tubes were determined using a portable WAL 0–4 bar absolute membrane pressure unit (Wal Mess-und Regelsysteme GmbH of Oldenburg, Germany).

H_2_, CO, CO_2_, CH_4_, and N_2_ contents were quantified using a Thermo Fisher Scientific Trace 1310 gas chromatograph with a thermal conductivity detector. In these determinations, 400 μL aliquots of gas were taken from the headspaces of culture tubes and were added to a packed Supelco Carboxen-1000 column (Gas Syringe A-2, Macherey-Nagel of Düren, Germany). The column was then heated to 108°C and maintained at this temperature for 6.25 min, followed by heating to 177°C at 120°C/min and maintenance at this temperature for 3.2 min. Finally, the column was heated at 120°C/min to 222°C and was maintained at this temperature for 4.6 min. Argon was used as the carrier gas at a flow rate of 11 mL/min. The injector and detector were adjusted to 125°C and 235°C, respectively. Measured GC values were standardized to 100%. Finally, methane production rates (MPR) were calculated as ml of CH_4_ produced/(mL medium *h).

## Results

### Co-culture

*Carboxydocella thermautotrophica, Carboxydocella sporoprducens* (Carbos), and cells from the three thermophilic rod-shaped methanogenic archaeal isolates (TRMC) proliferated well together in the newly designed medium without vitamin solution or yeast extract and under the growth conditions are presented in Table 2 (11), (12).

**Table 2.** *Growth conditions*

| Growth conditions | *Carboxydocella*<br>*thermautotrophica*<br>(Range; Optimum) | *Carboxydocella*<br>*sporoproducens*<br>(Range; Optimum) | TRMC (Culture<br>Conditions:<br>MicroPyros) | Conditions<br>used for the co-culture (This<br>study) |
| --- | --- | --- | --- | --- |
| T [°C] | 40–68; 58 | 50–70; 60 | 63 | 63 |
| pH | 6.5–7.6; 7 | 6.2–8.0; 6.8 | 6.5 | 6.5 |
| Metabolic<br>reaction | CO+H<sub>2</sub>O->CO<sub>2</sub>+H<sub>2</sub> | CO+H<sub>2</sub>O->CO<sub>2</sub>+H<sub>2</sub> | CO<sub>2</sub>+4H<sub>2</sub><br>->CH<sub>4</sub>+2H<sub>2</sub>O | This study |

### Metabolic properties of TRMC isolates

Mixtures of the three methanogenic archaeal isolates methanated CO_2_ and consumed H_2_ from the gas phase in the volumes shown in Fig. 1. However, CO conversion was not observed with either steam gasifier or air gasifier gas (Fig. 1).

**Fig. 1:**
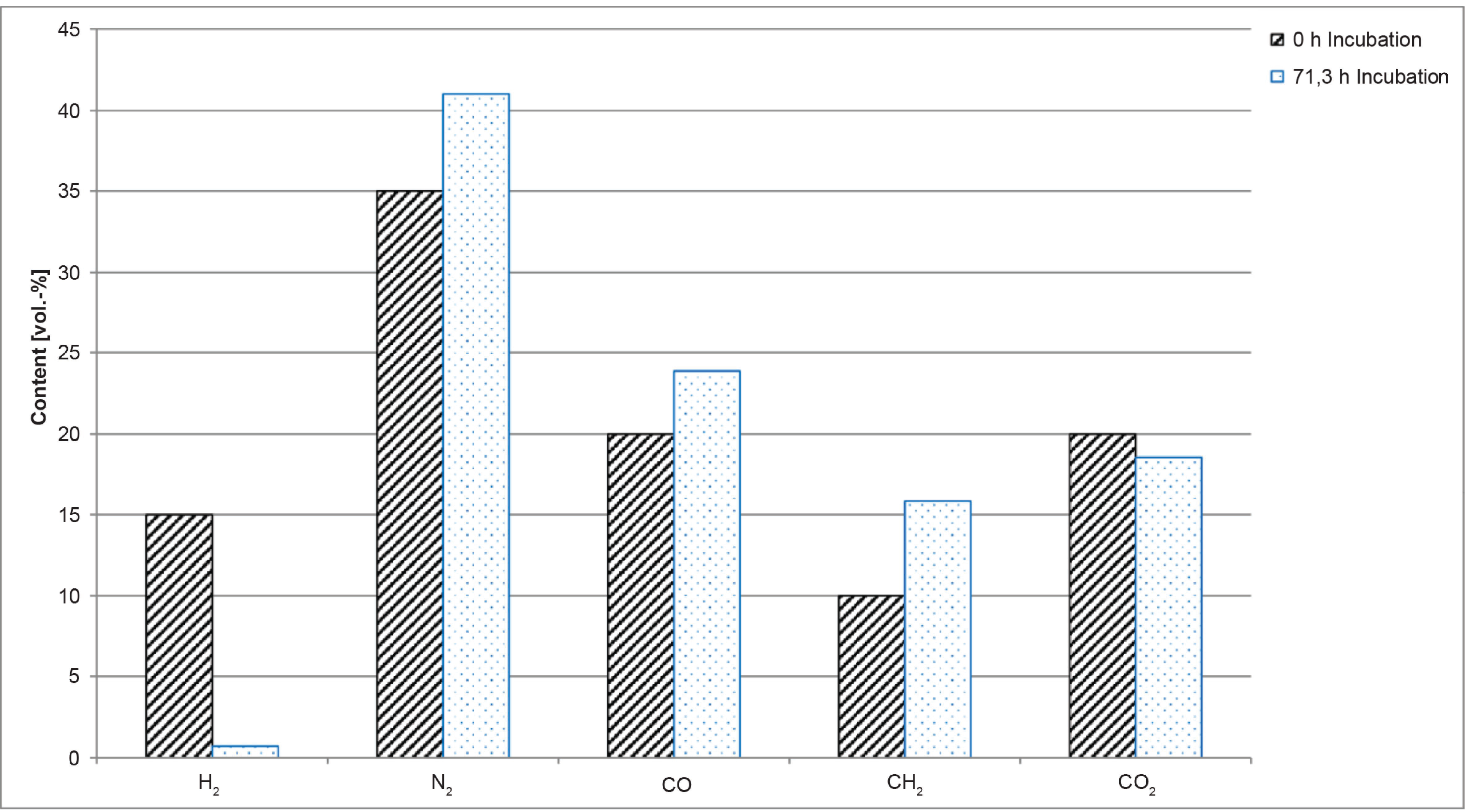
Gas compositions at 0 and 71.3-h incubation of TMRC in air gasifier gas

### Growth experiments with Carbos and TRMC

In initial experiments with combinations of Carbos and TRMC, microscope analyses showed the presence of fluorescent methanogenic archaea and the morphologically shorter non-fluorescent carboxydotrophic, hydrogenogenic bacteria growing in co-culture (Fig. 2), with an estimated TRMC to Carbos ratio of about 2:1 (Fig. 2).

**Fig. 2:**
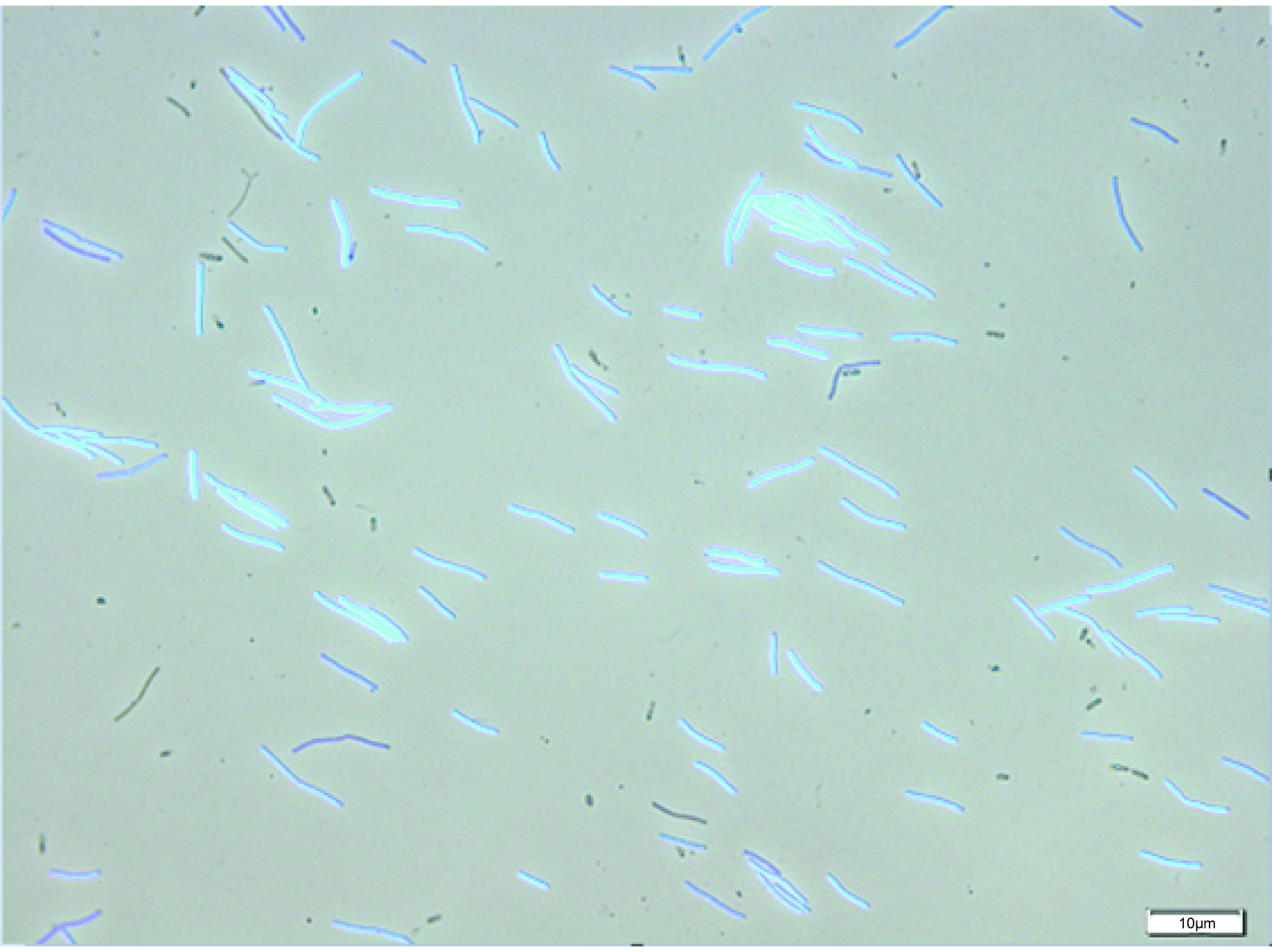
Co-culture of methanogenic archaea (TRMC; blue-colored rods) with carboxydotrophic bacteria (Carbos; black-colored rods) in steam gasifier gas after 96 h incubation at 63°C

### Consumption of steam gasifier gas by co-cultures

Figure 3 shows simultaneous decreases in H_2_ and CO concentrations and concomitant increases in CH_4_ contents of the gas phase. In these experiments, calculated maximum methane concentrations in the headspace were achieved within 96 h, based on reaction equations CO + H_2_O–> H_2_ + CO_2_ [1] and CO_2_ + 4 H_2_–> CH_4_ + 2 H_2_O [2]. Moreover, CO_2_ concentrations were increased immediately and then remained approximately constant, and pressure (p) profiles (p actual / p start) dropped during methanation to about 44% of the initial pressure.

**Fig. 3:**
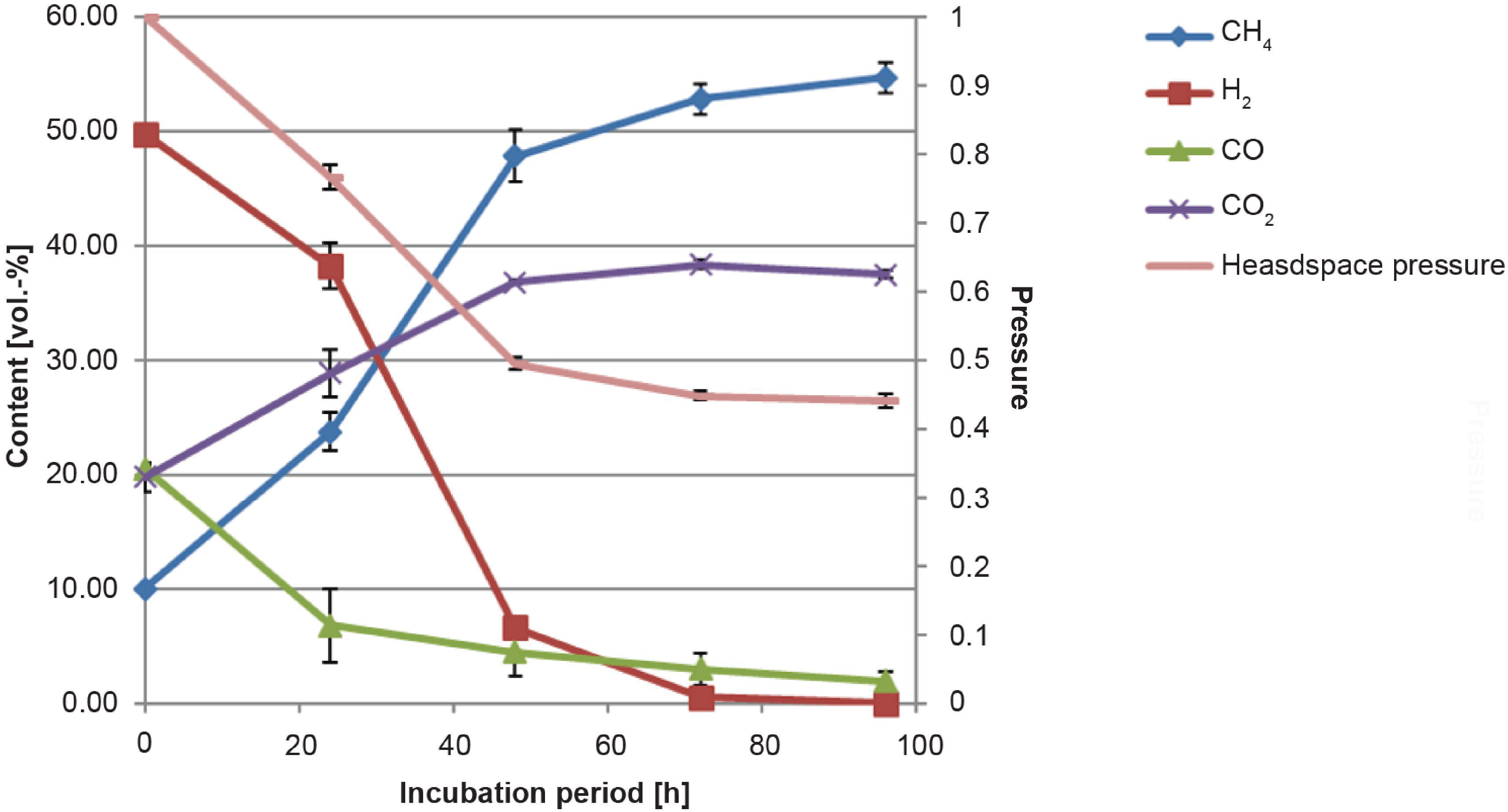
Changes in gas compositions and pressure during incubation in steam gasifier gas; error bars indicate standard deviations of three measurements

Initial hydrogen concentrations were 50% and dropped to 0.5% within 72 h, and CO concentrations decreased from 20% to 3% over the same time. Concomitantly, methane concentrations increased from 10% to 53%, and after 96 h incubation, H_2_ concentrations were 0%, CO concentrations were 2%, and methane concentrations were 55% of the entire volume. Moreover, increases in methane concentrations increased by about 1.0 vol%/h between 24 and 48 h, corresponding with a methane production rate of 0.03 mL/(mL*h).

The maximum possible methane concentration in the steam gasifier gas phase following complete H_2_ and CO methanation according to equations [1] and [2] was 55 vol%, and the total methane production rate from 10 vol% to 55 vol% methane over the entire 96 h incubation time was 0.02 mL/(mL*h).

During growth experiments in steam gasifier gas, numbers of cells in Carbos and TRMC increased during the first period of methanation, and then decreased and remained stable at about 1 × 10^8^ cells/mL (Fig. 4).

**Fig. 4:**
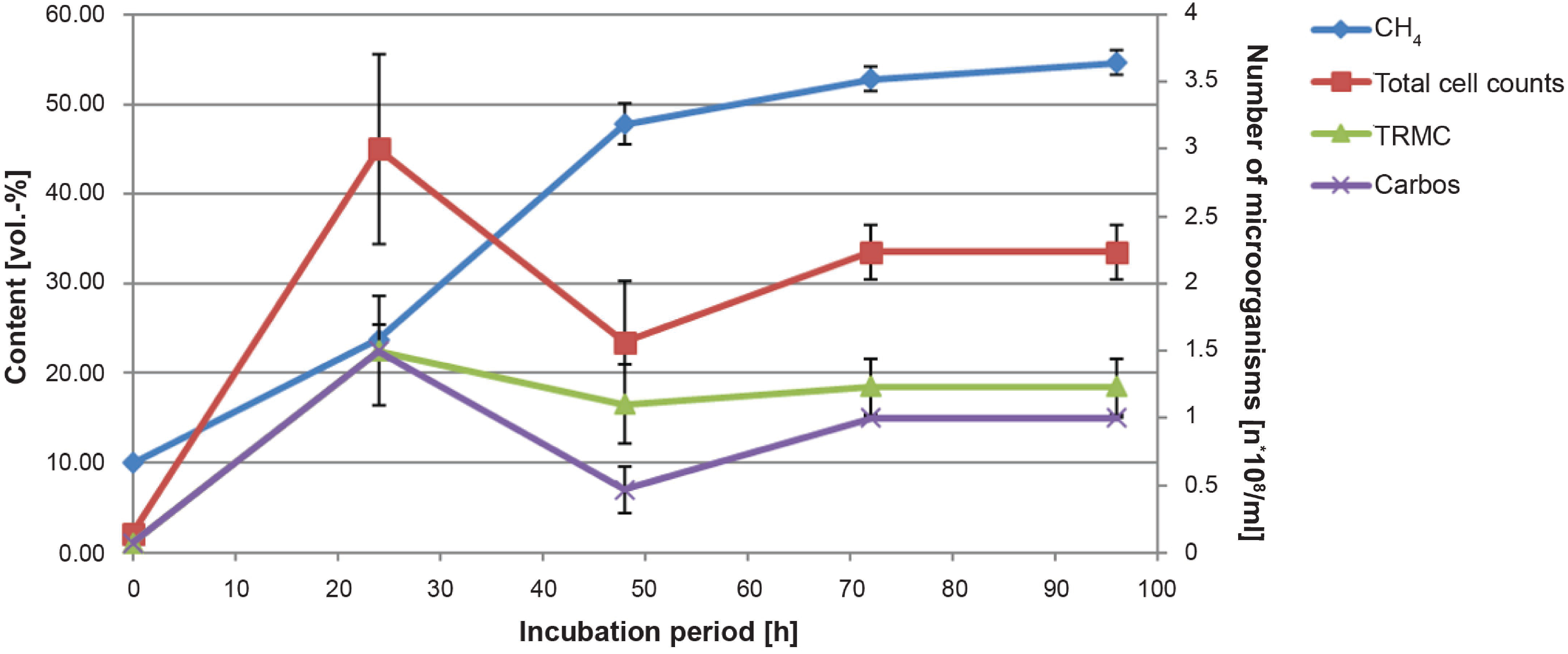
Changes in cell numbers over the incubation period in steam gasifier gas at 63°C; standard deviations of three measurements are indicated by error bars

### Consumption of air gasifier gas by co-cultures

H_2_ and CO contents decreased with increases in CH_4_ concentrations in the gas phase (Figure 5), and the calculated maximum methane concentration in the headspace was achieved within 72 h. Simultaneously, pressure profiles in the gas phase of the glass tubes (p actual / p start) dropped to about 69% of the initial pressure during methanation. In accordance, hydrogen concentrations in air gasifier gas dropped from 15% to 1.5% over 48 h. Concomitantly, CO concentrations decreased from 20% to 3%, and methane concentrations increased from 10% to 21%. After 96 h, the hydrogen concentration was 0%, the CO concentration was 1%, and the methane concentration was 22% of the total volume, and this concentration was maximal in air gasifier gas following complete H_2_ and CO methanation, according to equations [1] and [2]. Moreover, in calculations of MPR, methane concentrations increased by 0.22 vol%/h between 24 and 48 h incubation, corresponding with a production rate of 0.01 mL/(mL*h). Similarly, the total methane production rate from 10 vol% methane at 0 h to 22 vol% at 72 h was 0.01 mL/(mL*h).

**Fig. 5:**
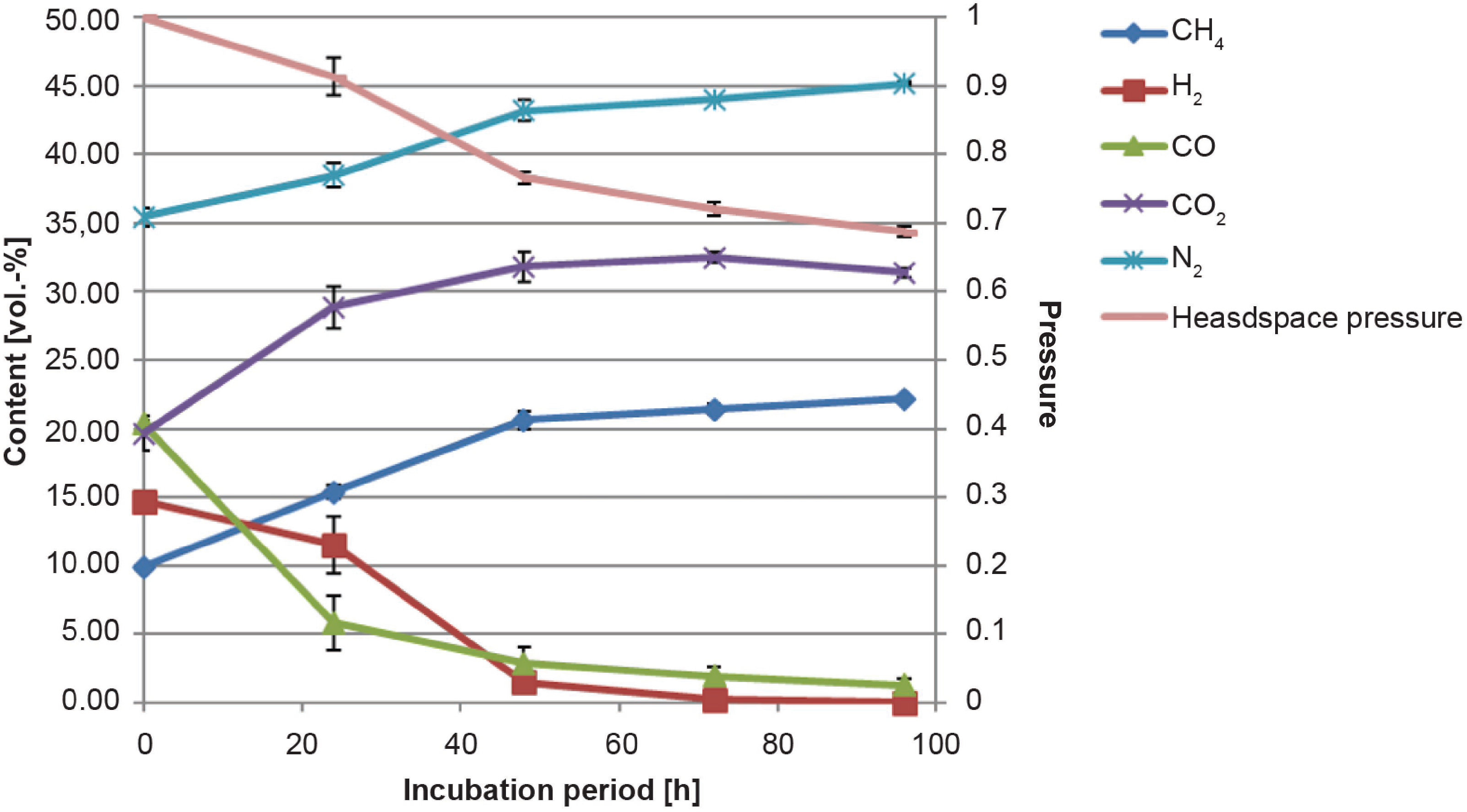
Changes in gas composition and pressure over the incubation period in air gasifier gas; standard deviations of three measurements are indicated by error bars

Growth experiments in air gasifier gas showed initial increases in cell numbers in Carbos and TRMC, followed by decreases to 0.5 × 10^8^/mL and 0.8 × 10^8^/mL, respectively (Fig. 6).

**Fig. 6:**
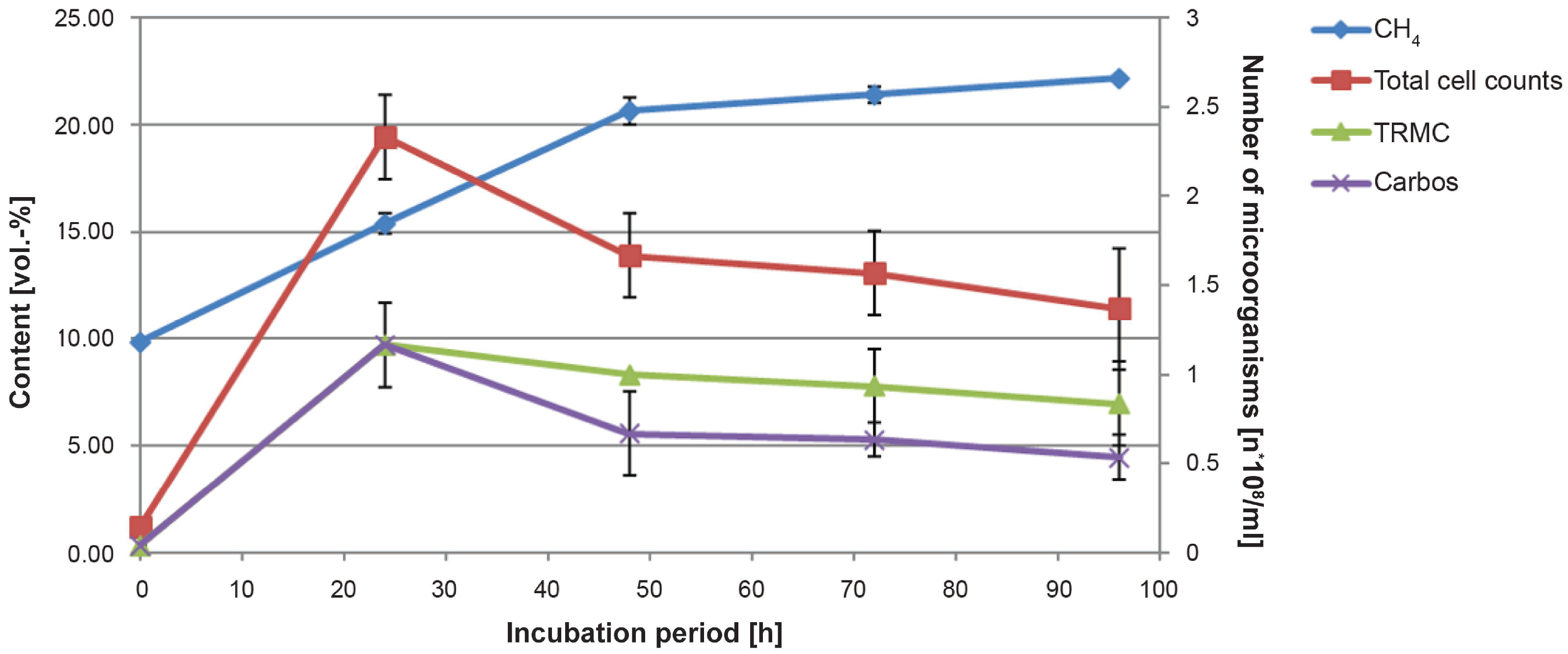
Changes cell numbers over the incubation period in air gasifier gas; standard deviations of three measurements are indicated by error bars

## Discussion

In the present novel combination cultures of thermophilic methanogenic archaea and thermophilic carboxydotrophic and hydrogenogenic bacteria, growth of all three organisms in synthetic steam gasifier gas and in synthetic air gasifier gas was observed, and CO, CO_2_, and H_2_ were converted into methane. In addition, the ensuing increases in methane concentrations corresponded with the stoichiometric ratios indicated by the equations CO + H_2_O -> H_2_ + CO_2_ and CO_2_ + 4 H_2_–> CH_4_ + 2 H_2_O.

Consistent with kinetic expectations, methane production slowed with decreasing reactant gas concentrations and lower pressures in culture tubes, reflecting decreased substrate gas availability. However, in air gasifier gas, methanation proceeded more slowly due to lower hydrogen partial pressures compared with those in steam gasifier gas. Klasson *et al.* previously showed simultaneous conversion of the gases CO and H_2_ by *Rhodospirillum rubrum* and H_2_ and CO_2_ by *Methanobacterium formicicum* and *Methanosarcina barkeri* in growing co-cultures at 34°C. In their study, *R. rubrum* growth was dependent on the presence of tungsten, light, and a carbon source other than CO, such as sugars, acetate, or yeast extract, and vitamin supplements were also provided (10), (17). In contrast, the present unique microbial combination grew optimally at 63°C and remained viable in the absence of light and expensive vitamin supplements and carbon sources, thus increasing the convenience and cost-effectiveness of large-scale cultures. Moreover, in the study by Klasson *et al.*, methane yields from H_2_ were 83% of the theoretical maximal yield (10), whereas we achieved 100% of theoretical maximal methane concentrations in the gas phase using synthetic steam and air gasifier gases. As a final advantage, the present co-culture system with spore-forming *Carboxydocella sporoproducens* was robust to periods of CO deficiency.

In summary, we demonstrated the use of a novel co-culture tool that efficiently allows the use of biowastes as sources of carbon and energy using gasification followed by biological methanation. This novel technology could also be applied to other CO_2_, H_2_, and CO-containing gases, such as exhaust gases from steel production, allowing use as raw materials for the production of CO_2_-neutral methane in industrial processes (18).

## Acknowledgments

We thank Dr. Jürgen Pettrak, Barbora Hemauer, Alexandra Fischer, and Maria Schubmann for their support in the lab and thank the whole team of the MicroPyros GmbH for fruitful discussions.

We also thank the SER Stadtentwässerung and Straßenreinigung Straubing for the permission to use their laboratory.

This work was supported by the TUM Applied Technology Forum Scholarship, so thank you, Christiane Hamacher.

## References

1. Ren J, Fedele A, Mason M, Manzardo A, Scipioni A. 2013. Fuzzy Multi-actor Multi-criteria Decision Making for sustainability assessment of biomass-based technologies for hydrogen production. Int J Hydrog Energy 38:9111–9120.

2. Redaktion Onetz OM-DN. Interview mit Prof. Dr. rer nat. Andreas Hornung: Biobatterie wandelt Energie in biogene …: Biobatterie aus der Oberpfalz. onetzde.

3. Bär K, Mörs F, Götz M, Graf F. 2015. Vergleich der biologischen und katalytischen Methanisierung für den Einsatz bei PtG-Konzepten. gwf - GasErdgas 156:466–473.

4. Daniels L, Fuchs G, Thauer RK, Zeikus JG. 1977. Carbon monoxide oxidation by methanogenic bacteria. J Bacteriol 132:118–126.

5. Liu Y, Whitman WB. 2008. Metabolic, phylogenetic, and ecological diversity of the methanogenic archaea. Ann N Y Acad Sci 1125:171–189.

6. Ferry JG. 2010. CO in methanogenesis. Ann Microbiol 60:1–12.

7. Diender M, Stams AJM, Sousa DZ. 2015. Pathways and bioenergetics of anaerobic carbon monoxide fermentation. Front Microbiol 6:1275.

8. Schwede S, Bruchmann F, Thorin E, Gerber M. 2017. Biological syngas methanation via immobilized methanogenic archaea on biochar. Energy Procedia 105:823–829.

9. Tiquia-Arashiro SM. 2014. CO-oxidizing microorganisms, p 11–28. In Thermophilic carboxydotrophs and their applications in biotechnology. Springer, Cham.

10. Klasson KT, Cowger JP, Ko CW, Vega JL, Clausen EC, Gaddy JL. 1990. Methane production from synthesis gas using a mixed culture ofR. rubrum M. barkeri, and M. formicicum. Appl Biochem Biotechnol 24–25:317–328.

11. Hofbauer H, Kaltschmitt M, Keil F, Neuling U, Wagner H. 2016. Vergasung in der Gasatmosphäre, p 1059–1182. In Energie aus Biomasse. Springer Vieweg, Berlin, Heidelberg.

12. Sokolova TG, Kostrikina NA, Chernyh NA, Tourova TP, Kolganova TV, Bonch-Osmolovskaya EA. 2002. Carboxydocella thermautotrophica gen. nov., sp. nov., a novel anaerobic, CO-utilizing thermophile from a Kamchatkan hot spring. Int J Syst Evol Microbiol 52:1961–1967.

13. Slepova TV, Sokolova TG, Lysenko AM, Tourova TP, Kolganova TV, Kamzolkina OV, Karpov GA, Bonch-Osmolovskaya EA. 2006. Carboxydocella sporoproducens sp. nov., a novel anaerobic CO-utilizing/H_2_-producing thermophilic bacterium from a Kamchatka hot spring. Int J Syst Evol Microbiol 56:797–800.

14. Zhao Y, Cimpoia R, Liu Z, Guiot SR. 2011. Orthogonal optimization of Carboxydothermus hydrogenoformans culture medium for hydrogen production from carbon monoxide by biological water-gas shift reaction. Int J Hydrog Energy 36:10655–10665.

15. Hungate RE. 1969. Chapter IV: A roll tube method for cultivation of strict anaerobes, p 117–132. In Norris JR, Ribbons DW (eds), Methods in microbiology. Academic Press.

16. Wolfe RS. 1971. Microbial formation of methane, p 107–146. In Rose AH, Wilkinson JF (eds), Advances in microbial physiology. Academic Press.

17. Klasson KT, Ackerson MD, Clausen EC, Gaddy JL. 1992. Bioconversion of synthesis gas into liquid or gaseous fuels. Enzyme Microb Technol 14:602–608.

18. Ahrens R. 2015. ThyssenKrupp sucht die große LösungVDI Nachrichten Nr. 10/2015.

